# PharmAverse: An Interactive Dashboard Using MedDRA Hierarchy

**DOI:** 10.1101/2025.06.17.660110

**Authors:** Abir Omran, Barbara Füzi, Alexander Amberg, Gerhard F. Ecker

## Abstract

The Medical Dictionary for Regulatory Activities (MedDRA) is a standardized international medical terminology widely used in databases that collect adverse event data for drugs. MedDRA employs a hierarchical structure, allowing for analysis at different levels of granularity. However, as the volume of adverse event data continues to grow, and given MedDRA’s complexity, it becomes increasingly challenging to analyze, and visualize the information effectively. To address this, PharmAverse was developed—an interactive dashboard designed to facilitate the analysis and visualization of adverse event data using MedDRA terms. It offers three main views: Adverse Events, Drug, and Target.

## Introduction

Various databases containing adverse event data for drugs, such as the FDA Adverse Event Reporting System (FAERS) and PharmaPendium, use a standardized international medical terminology known as the Medical Dictionary for Regulatory Activities (MedDRA) terminology [1], [2]. MedDRA serves as the ontology for adverse events and follows a hierarchical structure consisting of five levels. The highest level is the “System Organ Classes”, followed by “High Level Group Terms”, “High Level Terms”, “Preferred Terms” and finally, “Lowest Level Terms”. The Lowest Level Terms are the expressions used in practice to report observations. However, in databases such as FAERS, Preferred Terms are used for adverse event reporting. By utilizing the different hierarchical levels, one can better understand how the various adverse events are related. It also enables a more global examination of a drug’s adverse events by analyzing how the drug affects different system organs. Due to the hierarchical structure and depending on the dataset size, it can be challenging to manually analyze the data efficiently. Therefore, we developed PharmAverse, a dashboard that utilizes MedDRA’s hierarchy to visualize adverse event data through three views: Adverse Events, Drug, and Target. This tool is easily integrable and supports the use of preclinical, clinical, and post-marketing data. Furthermore, this tool can be used with adverse event data for any type of drug from any database that utilizes MedDRA terminology. Additionally, the plots can be downloaded for any downstream application. As a use case, we applied the dashboard to therapeutic antibody data to demonstrate its visualization capabilities.

## Features

### General Features

The dashboard is implemented in Python using Dash, a framework for building machine learning and data science web applications. The dashboard includes three views: Adverse Events, Drug, and Target. Each view features a dropdown menu that allows the user to select from the available data types—preclinical, clinical, or post-marketing—depending on what has been provided by the user. Additionally, the user can modify the plot size by selecting the desired plot through the dropdown menu on the right side and adjusting the width and height using the sliders under the dropdown menu. If a bar plot is selected, the font size of the x-axis labels can also be adjusted. All plots can be saved as PNG files by clicking the camera icon in the upper right corner of each plot. Figure 1 illustrates the Adverse Events view and highlights the features available across all views (in black) and the features unique to the Adverse Events view (in blue).

**Figure 1:**
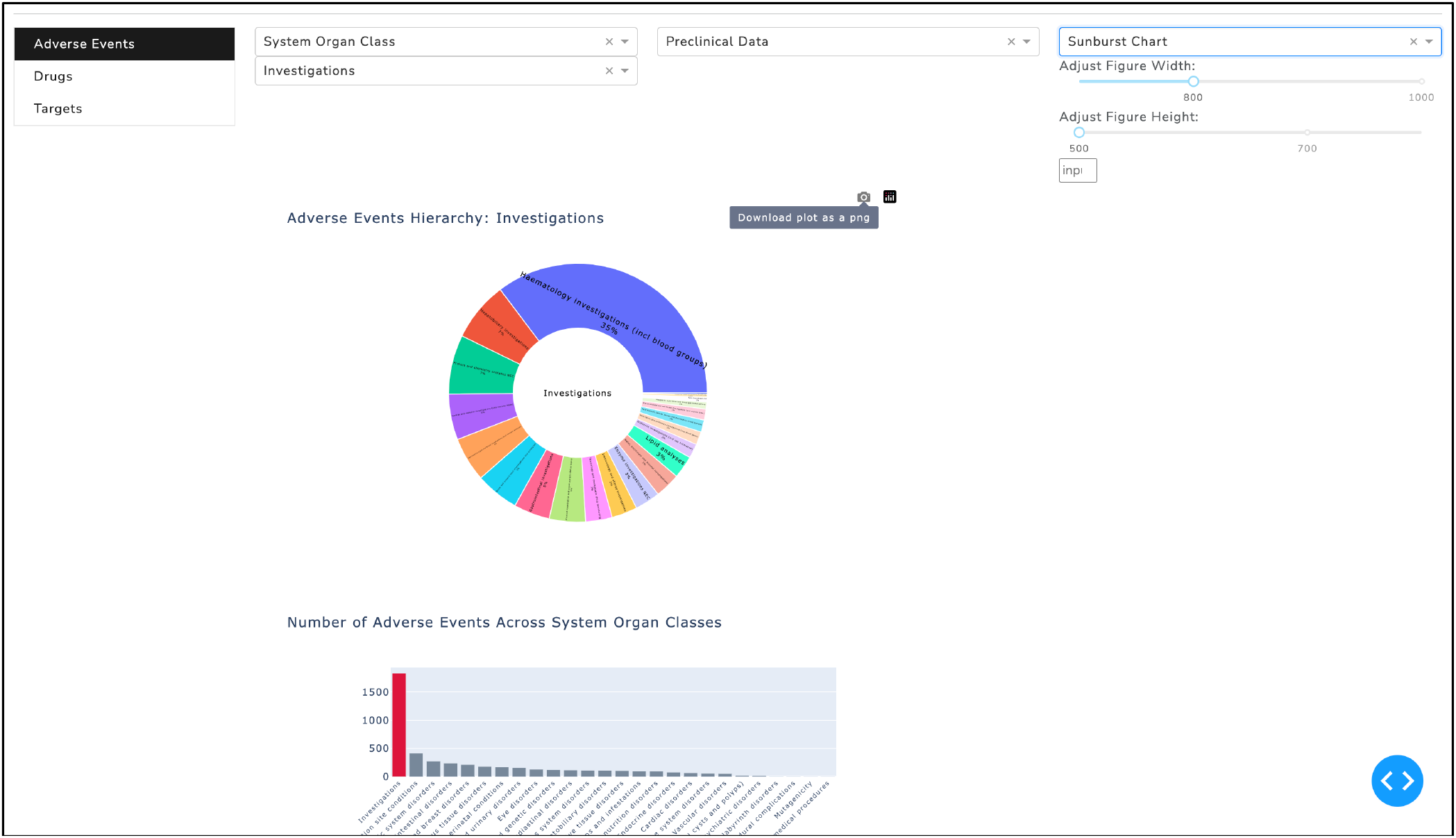
Overview of the Adverse Events view in the dashboard. Features available across all views are marked in black text, while features specific to the Adverse Events view are highlighted in blue. The figure also indicates which plots are interactive. The tabs on the left allow users to navigate between the available views.

### Adverse Events View

The Adverse Events view is the initial view displayed upon opening the dashboard — it enables users to analyze the data according to the hierarchical levels defined by MedDRA. This view includes three interactive plots: The first plot is a sunburst chart that displays lower-level adverse event terms linked to the selected System Organ Class from the dropdown menu (see Figure 1). The second plot is a bar chart that visualizes the count of Preferred Terms for each System Organ Class, highlighted only the selected class. Users can interact with the sunburst chart by selecting an adverse event term, which will then update both pie chart and the bar chart. If preclinical data is used, a pie chart appears showing the species that reported Preferred Terms associated with the selected adverse event term. In contrast, if clinical or post-marketing data is used, a bar chart will be displayed, which visualizes all drugs reporting the selected adverse event term.

### Drug View

The Drug view enables the user to analyze data based on the selected drug. Depending on the available study data, up to four plots can be displayed in this view. The interactive bar chart shows the count of Preferred Terms for each System Organ Class associated with the selected drug. The user can select a System Organ Class from the bar chart, which will update the sunburst chart to display its lower-level terms. Additionally, the sunburst chart is interactive, allowing the user to select an adverse event term at any level to visualize the species in the pie chart which is reporting the adverse events for the selected term when preclinical data is displayed. The pie chart allows the user to select a species, updating the bar chart above to show the count of adverse events (in Preferred Terms) associated with the selected species.When clinical or post-marketing data is used, the bar chart visualizes the counts of Preferred Terms for all drugs reporting the adverse event terms selected from the sunburst chart.

In Supplementary Figure 1, the adverse events for adalimumab are visualized using preclinical data. The System Organ Classes are presented in the bar chart, with the counts for each class displayed. By selecting a System Organ Class, in this case, “Investigation”, the adverse events related to that class are shown in a hierarchical manner in the sunburst chart. Since preclinical data is selected, the pie chart shows the proportion of species reporting the adverse event selected from the sunburst chart, in this case, “White blood cell analyses”.Selecting a species from the pie chart updates the bar chart to display the adverse events associated with the chosen species for all drugs and adalimumab, which is the selected drug is highlighted in red.

### Target View

The Target view allows the user to analyze data based on the selected target. This view includes three available plots. By selecting a target from the dropdown menu, the sunburst chart displays the drug(s) associated with the chosen target and indicates whether the drug(s) are linked to other targets. The bar chart visualizes the count of adverse events (in Preferred Terms) for the selected target across all System Organ Classes. Additionally, the user can select a System Organ Class from the bar chart, which updates the second sunburst chart. The second sunburst chart then visualizes the lower-level adverse event terms associated with the chosen System Organ Class.

The Supplementary Figure 2 illustrates the Target view for Tumor Necrosis Factor-alpha (TNF-α). The upper sunburst chart displays all the TNF-α-targeting drugs included in the dataset. The interactive bar plot shows the adverse events counts reported for these drugs across all System Organ Classes. By selecting the System Organ Class “Infections and infestations” from the bar plot, the lower sunburst chart will update to display the MedDRA term hierarchy corresponding to the “Infections and infestations”.

### Data Structure and Requirements

A MedDRA license is required to access the MedDRA release package. Begin by downloading the MedDRA release package and place the MedAscii folder in the working directory, this folder contains the MedDRA terminology. The command below generates a dataset that maps MedDRA terms to their respective hierarchical levels. The resulting file will then be used in the data processing step.

`python meddra_hierarchy_mapping.py`

To prepare the data for use in the dashboard, ensure that the dataset contains two columns: the first column should represent the drug (e.g., adalimumab), and the second should contain the preferred terms according to MedDRA (e.g., Spleen disorder). For preclinical data, the third column should include species information (e.g., Cynomolgus monkey). The following command processes the data, saves it to the designated folder, and prepares it for integration into the dashboard.

`python data_processing.py –-file_path <path_to_the_file.txt> -- study_type <preclinical | clinical | postmarketing>`

Additionally, for the Targets view, an extra dataset is required, containing drugs in the first column and their corresponding targets in the second column. This target dataset should be placed in the datasets folder under the name “target_data.csv”.

### Availability and Implementation

The project folder containing the source code is available here: https://github.com/PharminfoVienna/PharmAverse

To launch the dashboard, navigate to the project folder and run the following command in the terminal:

`python app.py`

Then, open your web browser and go to the following address: http://127.0.0.1:8050/

This will launch the dashboard locally on your machine.

## Acknowledgements

In this work the MedDRA® terminology used which is the international medical terminology developed under the auspices of the International Council for Harmonisation of Technical Requirements for Pharmaceuticals for Human Use (ICH).

## Funding

This research was funded in whole or in part by the Austrian Science Fund (FWF) W1232. For open access purposes, the author has applied a CC BY public copyright license to any author accepted manuscript version arising from this submission. The study was also funded by Sanofi.

## Notes

### Competing Interest Statement

The authors have declared no competing interest.

### Summary of Updates

An acknowledgements section has been added

