## Supplementary Information for "PharmAverse: An Interactive Dashboard Using MedDRA Hierarchy"

### Supplementary Figures

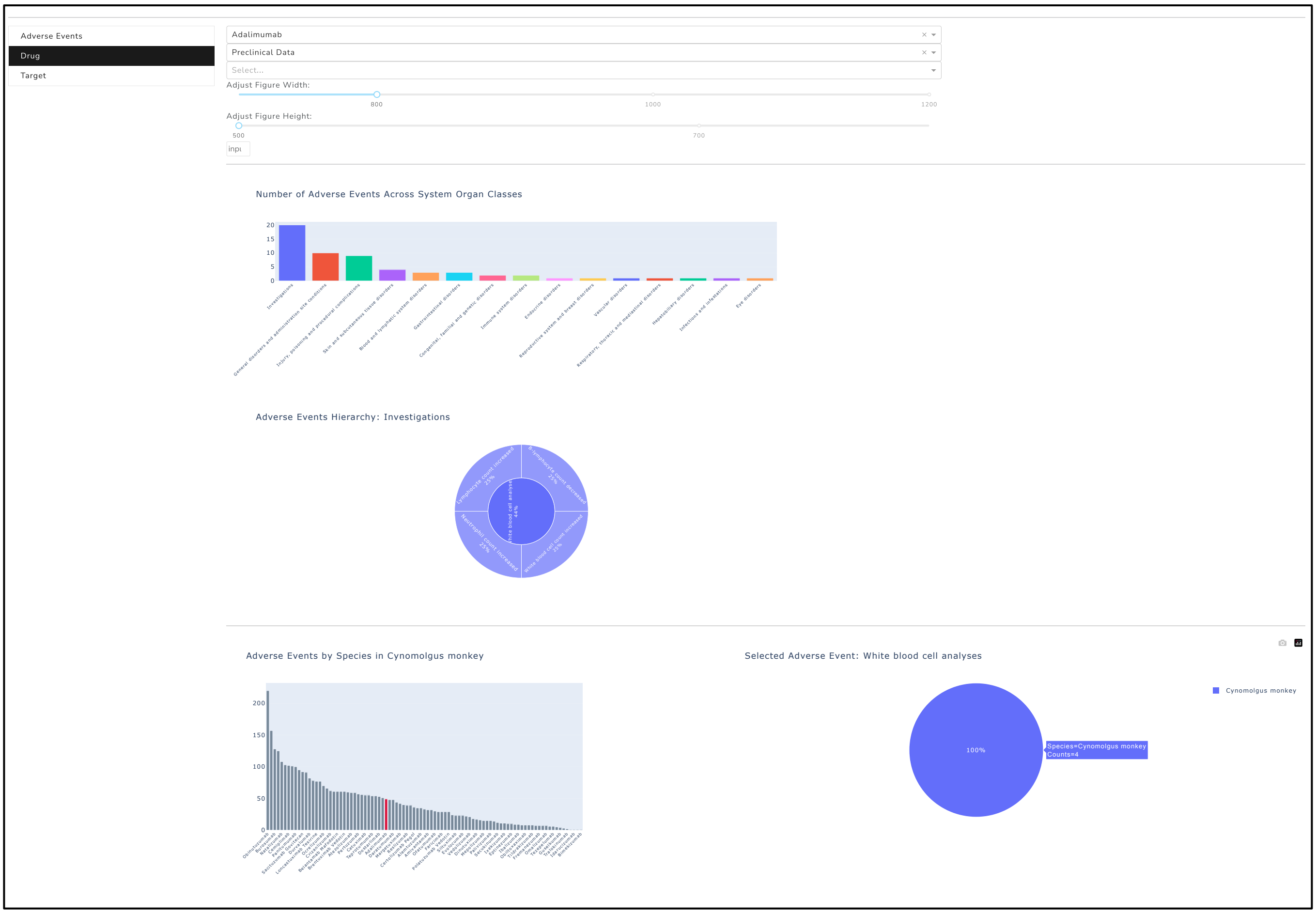


Supplementary Figure 1: The Drug view, where preclinical data for the selected drug, Adalimumab, is visualized. The interactive bar plot displaying the System Organ Classes allows the user to select categories, in this case "Investigation" was selected, which then shows the corresponding MedDRA terms in the Sunburst chart. When the adverse event "White blood cell analysis" is selected from the sunburst chart, it updates the pie plot to display the species reporting this adverse event. By clicking on the species displayed in the pie plot, the bar plot shows the adverse event count for all drugs reporting the chosen species.

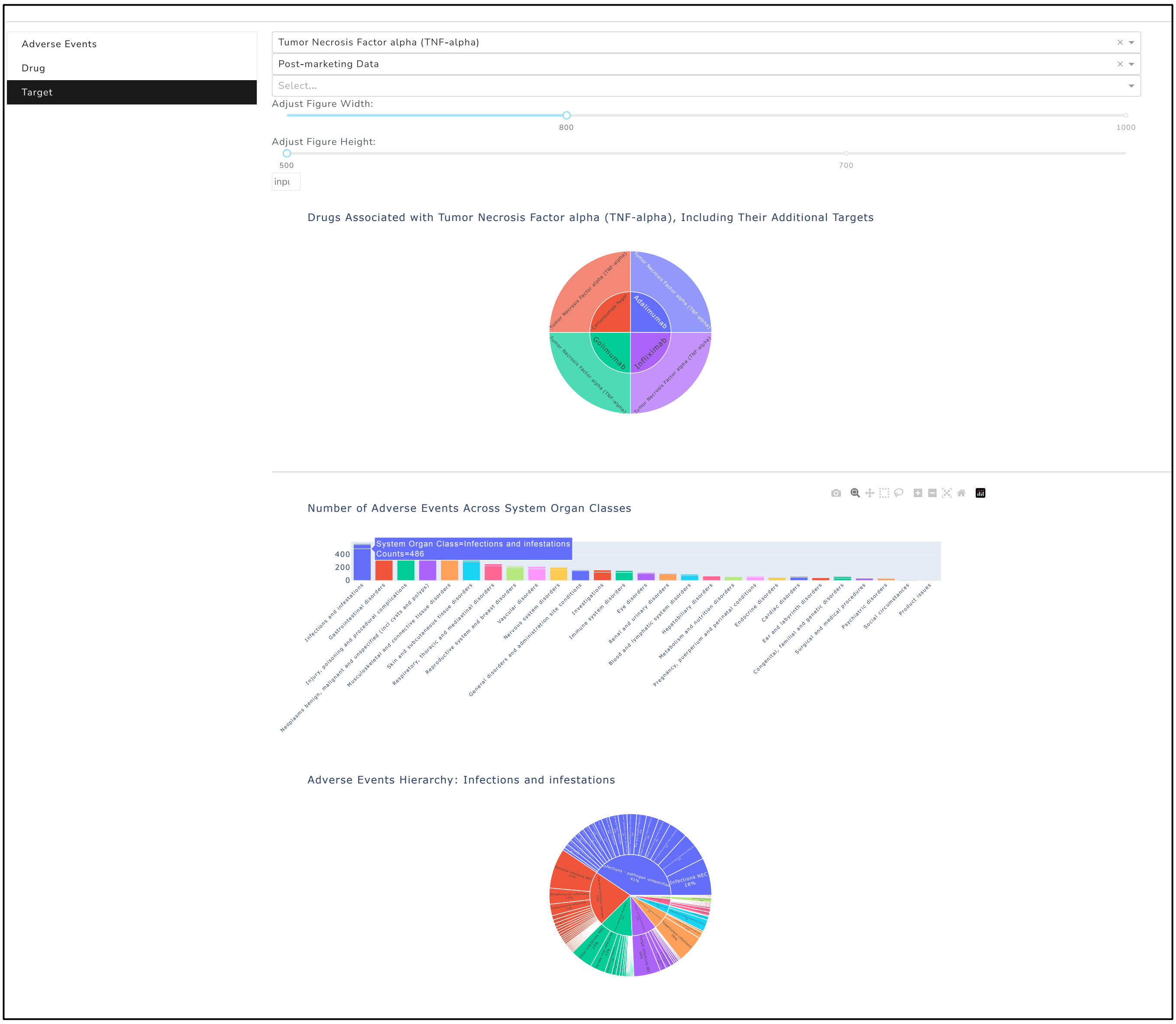


Supplementary Figure 2: The Target view for Tumor Necrosis Factor alpha (TNF-α) is selected as the target is visualized. The upper sunburst chart displays the drugs targeting TNF-α, while the bar plot shows the adverse event count across System Organ Classes for the selected target. By selecting the System Organ Class “Infections and infestations” from the bar plot, the lower sunburst chart displays the corresponding MedDRA terms.
